# Computational prediction of Drug-Disease association based on Graph-regularized one bit Matrix completion

**DOI:** 10.1101/2020.04.02.020891

**Authors:** Aanchal Mongia, Emilie Chouzenoux, Angshul Majumdar

## Abstract

**Motivation:** Investigation of existing drugs is an effective alternative to discovery of new drugs for treating diseases. This task of drug re-positioning can be assisted by various kinds of computational methods to predict the best indication for a drug given the open-source biological datasets. Owing to the fact that similar drugs tend to have common pathways and disease indications, the association matrix is assumed to be of low-rank structure. Hence, the problem of drug-disease association prediction can been modelled as a low-rank matrix-completion problem.

**Results:** In this work, we propose a novel matrix completion framework which makes use of the side-information associated with drugs/diseases for the prediction of drug-disease indications modelled as neighborhood graph: Graph regularized 1-bit matrix compeltion (GR1BMC). The algorithm is specially designed for binary data and uses parallel proximal algorithm to solve the aforesaid minimization problem taking into account all the constraints including the neighborhood graph incorporation and restricting predicted scores within the specified range. The results of the proposed algorithm have been validated on two standard drug-disease association databases (Fdataset and Cdataset) by evaluating the AUC across the 10-fold cross validation splits. The usage of the method is also evaluated through a case study where top 5 indications are predicted for novel drugs and diseases, which then are verified with the CTD database. The results of these experiments demonstrate the practical usage and superiority of the proposed approach over the benchmark methods.

## 1 Introduction

Inspite of the large financial investment in pharmaceutical industry, the number of drugs approved over the past few decades is limited Walters *et al.* (2011). This can be attributed to the time (10-15 years) and effort it takes to test a therapeutic compounds and declare it as a market-ready drug. The problem calls for an alternative to drug discovery: “drug-repositioning” or “drug-repurposing”. This essentially means that an existing, already approved drug is identified to seek for its new indications. The benefits include shorter drug-development timelines, established safety and savings on money for launching the drug. Also, the strategy of drug-repurposing offers an opportunity to overcome the threats associated with antimicrobial resistance (AMR) Kaul *et al.* (2019). Some examples of re-positioned drugs include chlorocyclizine, an anti-allergic drug re-purposed as an antiviral He *et al.* (2015), sertraline, an antidepressant drug as an antifungal Villanueva-Lozano *et al.* (2018) and disulfiram, an anti-alcoholic drug repurposed as an antibacterial Das *et al.* (2019); Thakare *et al.* (2019).

There have been some successfully re-positioned drugs through manual and rational investigations but this is not an efficient and scalable way given the huge space of drug interactions. Therefore, computational approaches have been used over the past years to systematically predict the indications, pruning down the massive search space for researchers and saving huge amounts of efforts, time and cost. This explains the immense importance of predicting new associations between drugs and diseases using statistical and machine learning based methods.

Initial attempts to predict novel indications were based on gene expression profiles Lamb *et al.* (2006); Hu and Agarwal (2009). Lamb *et al.* (2006) proposed a database having ranked drug response gene expression which were queried with a gene signature specific to a disease. The drug response profiles which either correlate or anti-correlate were identified. This approach lacks validation on a large scale dataset and may not be precise enough owing to different conditions under which expression profiles are generated.

Other set of approaches captured the notion of similarity Chiang and Butte (2009) where it was assumed that alternative for one of the two diseases which are treated by the same drug, may also be used as a potential treatment for the other disease.

Later, network-based models were proposed. Gottlieb *et al.* (2011) proposed PREDICT, a method which computationally predicts drug-disease associations using integrated drug and disease information. Various kinds of drug and disease similarities are calculated to find the feature vectors for the candidate associations which are further used to train a classification model using logistic regression. Wang *et al.* (2014) created a 3-layer heterogeneous network, corresponding to drug, disease and targets. Edge weights between the nodes of same type (i.e. intra-connections) correspond to similarity between them while those between different types of nodes are associated with the relationship or association between the nodes i.e. drug-target or drug-disease relationship. The missing edges of this network are inferred using guilt-by-association principle. In a similar fashion, Martinez *et al.* (2015) integrated information from drugs, diseases and targets and proposed a network-based prioritization method for predicting new drug indications and novel disease treatments. Another work, Wang *et al.* (2013) integrates molecular structure, molecular activity, and phenotype data and constructs a kernel function to correlate drugs with diseases, and finally train an SVM (Support vector machine) classifier for the prediction of drug-disease interaction. Yu *et al.* (2016) identifies the drug/disease modules by clustering the drug network and disease network and then connecting drug-disease module pairs. Very recently, a new network-based approach was proposed by Yang at al Yang *et al.* (2019a) where the authors employ heterogeneous network embedding for the characterization of drug-disease association and trains an SVM for predicting novel associations.

There have also been several machine learning and deep learning techniques used for association prediction apart from the ones (clustering and classification methods) used in few of the works mentioned above. Very recently, Xuan *et al.* (2019) trained a dual convolutional neural network on two association layers simultaneously, one encoding the drug-disease characteristics while another one, the associated neighborhood information. Jiang *et al.* (2019) extracted feature descriptors from drug and disease Gaussian interaction profile based and other similarities using autoencoder and trained a random forest classifier to predict drug-disease associations. Wang *et al.* (2019) trained a neural network on the aggregated neighborhood information with the drugs and diseases association and similarity matrices; they minimize the loss between initial and recovered matrices while training the neural network on the heterogeneous data.

Drug-disease association prediction can also be modelled intuitively as a collaborative filtering problem. The objective of this class of approaches is to recover a complete matrix from its sampled entries by exploiting its low-rank structure. The low-rank assumption stems from the idea that similar drugs affect biological systems in a similar way and have common indications Jadamba and Shin (2016).

The underlying techniques which aim to solve collaborative filtering problem via matrix completion are majorly based on matrix factorization or nuclear norm minimization. Matrix factorization has been employed in the community over the past few years. It assumes that the number of latent (or hidden) features which may determine the association between a drug and a disease (such as substructures, targets, enzymes, pathways, MeSH information, etc) is very few and highly correlated. Yang *et al.* (2014) used probabilistic matrix factorization on causal networks connecting drug–target–pathway–gene–disease to classify drug-disease associations. Dai *et al.* (2015) integrates genomic space into the matrix factorization framework to exploit the molecular biological information using gene interaction network and then predicts novel indications. Zhang *et al.* (2018) projects the association information to two low-rank latent spaces, while taking into account the topological information of drug and disease data points by using the similarity information of drugs and diseases in the objective function of matrix factorization.Matrix factorization is a bilinear non-convex problem, which makes it challenging to solve, as spurious local minima usually occur. This problem can be overcome by an alternate approach for matrix completion: Nuclear norm minimization Candès and Recht (2009). Minimizing the nuclear norm (sum of singular values of a matrix) is the closest convex surrogate to minimizing the rank (number of singular values of a matrix) of that matrix, which is known to be a NP-hard problem. There are relatively few works modelling the prediction task using nuclear norm minimization. Luo *et al.* (2018) and Yang *et al.* (2019b) deploy nuclear norm minimization on a heterogeneous network matrix obtained by integrating drug similarity, disease similarity, association matrix and its transpose; the latter work additionally handles the noise originating from similarities which violate the low-rankness and restrict the predicted values to be in range [0,1]. But, the low-rank property of the heterogeneous matrix is unexplained in both the works, although it is a crucial assumption behind nuclear norm minimization. This heterogeneous matrix comprises of associations between drugs and diseases as well as drug-drug and disease-disease similarities. The authors clearly explain validity of the low-rank assumption in association matrix but not for the heterogeneous matrix.

In this work, we formulate drug disease association prediction as a one-bit matrix completion problem. Furthermore, we introduce graph regularization to exploit the similarities between drugs and diseases. The objective function is minimized using parallel proximal algorithm (PPXA) Pustelnik *et al.* (2011). PPXA is an iterative proximal splitting algorithm that paralelly solves for each of the non-necessarily smooth terms in the objective function, while benefiting from sounded convergence guarantees. The novelty of our approach lies in

- Modelling the drug-disease association prediction as graph-regularized matrix completion problem.
- Restricting the association scores in range [0,1] for obtaining meaningful biological scores.
- Solving the optimization problem using PPXA which has guaranteed convergence properties Pustelnik *et al.* (2011).

A schematic overview of GR1BMC is shown in Figure 1.

**Fig. 1.**
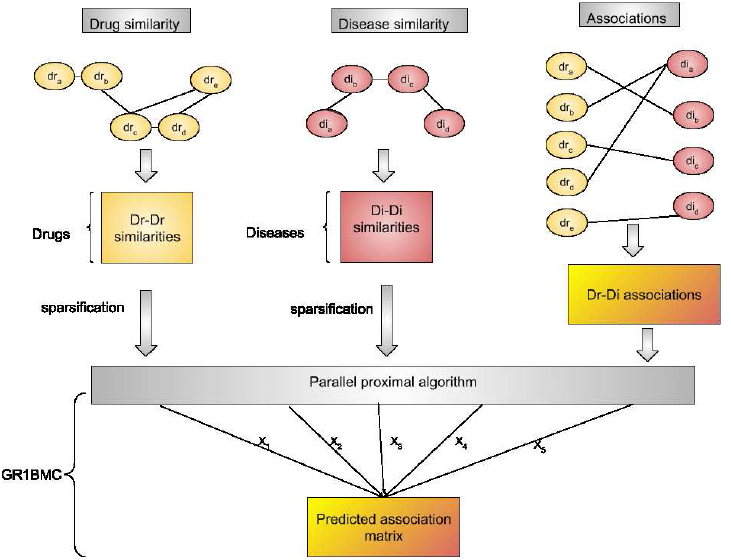
A schematic overview of GR1BMC for predicting drug-disease assocations

## 2 Material and Methods

### 2.1 Dataset

We have used two gold standard databases to validate our approach. The first one, called *F dataset*, proposed by Gottlieb *et al.* (2011) has 313 diseases, 593 drugs and 1933 drug-disease associations from various sources. The second dataset, called *Cdataset* is a larger one with 663 drugs, 409 diseases and 2532 associations Luo *et al.* (2016).

For the datasets the drug information is obtained from DrugBank Wishart *et al.* (2006), an exhaustive database containing comprehensive information about drugs and targets. The disease information was assembled from human phenotypes listed in public database, OMIM (Online Mendelian Inheritance in Man) database Hamosh *et al.* (2002), which has information on human genes and diseases.

The similarity information of drugs, calculated as Tanimato score Tanimoto (1958), is extracted using Chemical Development Kit (CDK) Steinbeck *et al.* (2003) based on the chemical structures of drugs in SMILES (Simplified Molecular-Input Line-Entry System) format, obtained from DrugBank. MimMiner Van Driel *et al.* (2006) provides the similarities between diseases using the medical descriptors of diseases from OMIM database by measuring the number of MeSH (medical subject headings vocabulary) terms. Both kinds of similarites are in range [0,1]. The information on number of drugs, diseases and the associations between them has been summarized in Table 1.

**Table 1.** A summary of the number of associations, drugs and diseases in each dataset used.

| Datasets | # Associations | # Drugs | # Diseases |
| --- | --- | --- | --- |
| # Fdataset | 1933 | 593 | 313 |
| # Cdataset | 2532 | 663 | 409 |

### 2.2 Pre-processing

We perform the following two steps to ensure better learning:

- SIMILARITY SPARSIFICATION: To ensure that the local geometries of the association data are preserved, both types of similarities are sparsified by keeping only *p*–nearest neighbor of each drug/disease profile in the drug/disease similarity matrix. This is done by elementwise multiplying the similarity matrix with another “neighborhood matrix” representing *p*-nearest neighbor graph of drugs/diseases.

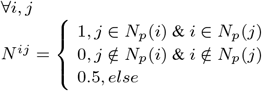

where *N_p_* (*i*) is the set of *p* nearest neighbors to drug *dr_i_*. We have set *p* =5 here.
- NORMALIZATION OF GRAPH LAPLACIANS: Instead of graph laplacians *L_di_* and *L_dr_*, we use their normalized versions 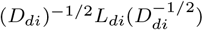 and (*D_dr_*)^-1/2^ *L_dr_* (*D_dr_*)^-1/2^ since they are known to perform better than their un-normalized counterparts Belkin *et al.* (2006)

### 2.3 Proposed Algorithm

To predict the drug-disease association matrix *X*, we model it as a low-rank matrix and aim to complete its available version *Y*. Since low-rank approximation is an NP-hard problem, we solve its closest convex surrogate i.e. nuclear norm minimization. Nuclear norm is defined as the sum of absolute singular values of a matrix.

To incorporate the disease and drug similarities into this imputation framework, we introduce the Laplacian graph regularization terms.

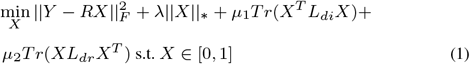

Here, ||·||* denotes the nuclear norm, *T_r_* denotes the trace, *R* is the binary masking operator which is element-wise multiplied to complete data matrix *X* (having 1’s at train indices and 0’s at test indices) and *L_di_* and *L_dr_* denote the disease and drug Laplacian matrices.

Here, we propose to make use of the parallel proximal algorithm (PPXA) from Pustelnik *et al.* (2011) (see also Cherni *et al.* (2017) for its application in the context of biochemistry). In this algorithm, we solve for *X*, by taking a proxy variable for each of the terms in(1) Combettes and Pesquet (2011) and an extra proxy variable *X*_3_ to ensure that the predicted scores are in range [0,1]. For each iteration k, we need to compute the following proximity operators:

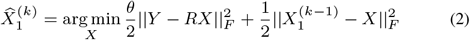

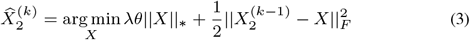

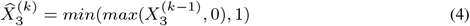

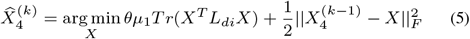

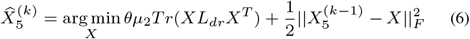

Hereabove, *θ* corresponds to the number of terms treated in parallel, that is *θ* = 5. Below we provide the solution of each of the above subproblems:

- Solving for 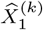 involves taking the gradient of (2) and equating to 0:

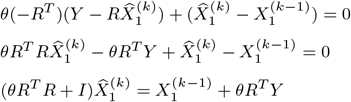

where I is the identity matrix. The above can now be easily solved by finding least squares solution.
- 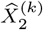 can be obtained by soft-thresholding the singular values of 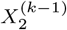 and multiplying the thresholded singular value matrix by the left and right singular vector matrices of 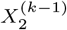 i.e.

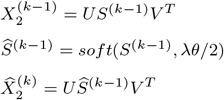

where *soft*(*S*^(*k*-1)^,λ*θ*/2) = *sign*(*S*^(*k*-1)^)*max*(0,|*S*^(*k*-1)^| − λ*θ*/2). Here *S*^(*k*-1)^ denotes the singular value matrix, *U* and *V* are the left and right singular matrices of 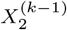 obtained after SVD-decomposition.
- Solving for 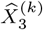 is done by applying max-thresholding followed by min-thresholding on 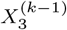.
- To solve for 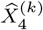, we employ the same strategy as for 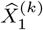 and equate the gradient of (5) to 0:

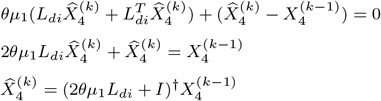
- Similarly, update step for 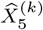 can be obtained as follows:

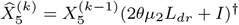

In the above two update steps, *A*^†^ denotes the Moore-Penrose pseudo inverse of A. The next iterate *X*^(*κ*)^ is finally obtained by averaging over the five proximal values, as follows:

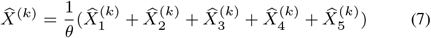

with *θ* = 5. Furthermore, each of the proxy variables is updated via the following update rule:

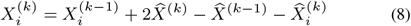

The complete algorithm is given in Algorithm 1^1^.

We display in Figures 2 and 3 example of convergence plots (i.e. evolution of objective function along iterations) for Fdataset and Cdataset, respectively.

**Fig. 2.**
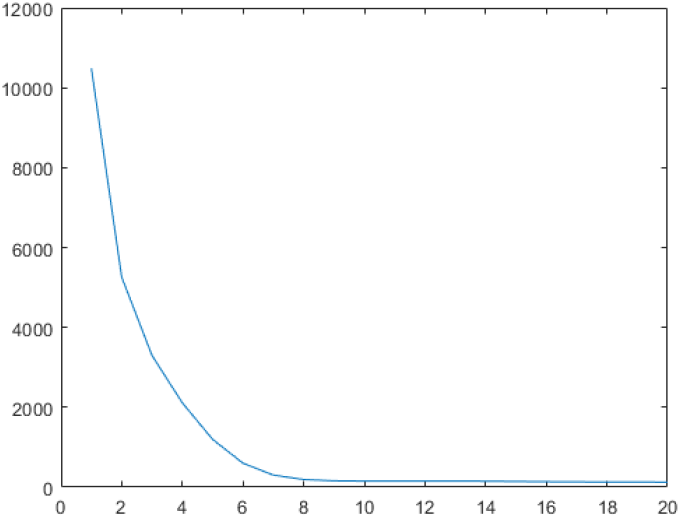
Convergence plot for GR1BMC on Fdataset

**Fig. 3.**
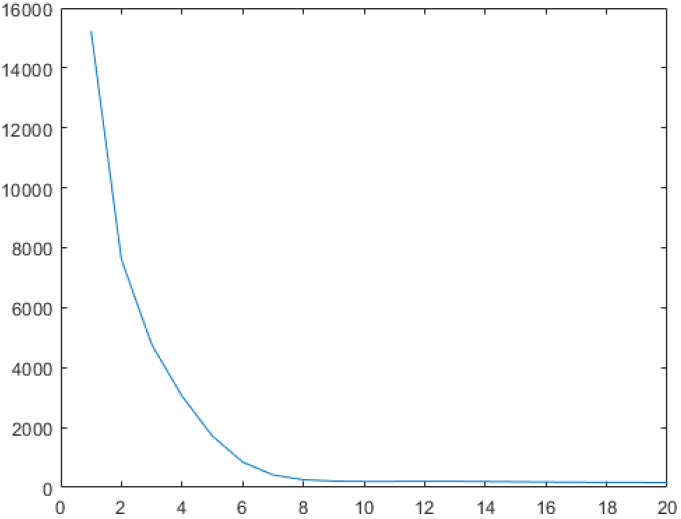
Converegence plot for GR1BMC on C dataset

## 3 Results and Discussion

### 3.1 Evaluation criteria

To experimentally evaluate the prediction performance of GR1BMC, we use *κ*-fold cross validation strategy (*κ* = 10). *κ*-fold cross validation is an evaluation method where we divide the known associations into *κ* equal subsets (called folds). Out of all the subsets, 1 of them is treated as a testing set, while the remaining ones constitute the training set. The associations in training set are given as input to the algorithm which then returns the fully imputed association matrix.

After this matrix completion, the predictions on testing set and other candidate associations for all drugs are ranked in descending order of scores and TPR (True Positive Rate)/Recall, FPR (False Positive Rate) and PPV (Positive predicted value)/Precision is calculated for every rank threshold. These values at every threshold are used to plot an ROC (Receiver Operating Characteristic) curve with FPR on x-axis and TPR on y-axis. In a similar way, a Precision-Recall curve is obtained by plotting Recall/TPR on x-axis and Precision on y-axis. The area under both these curves called Area under the ROC curve (AUC) and Area under the precision-recall curve (AUPR) are used to asses the performance of the methods used to predict drug-disease associations.

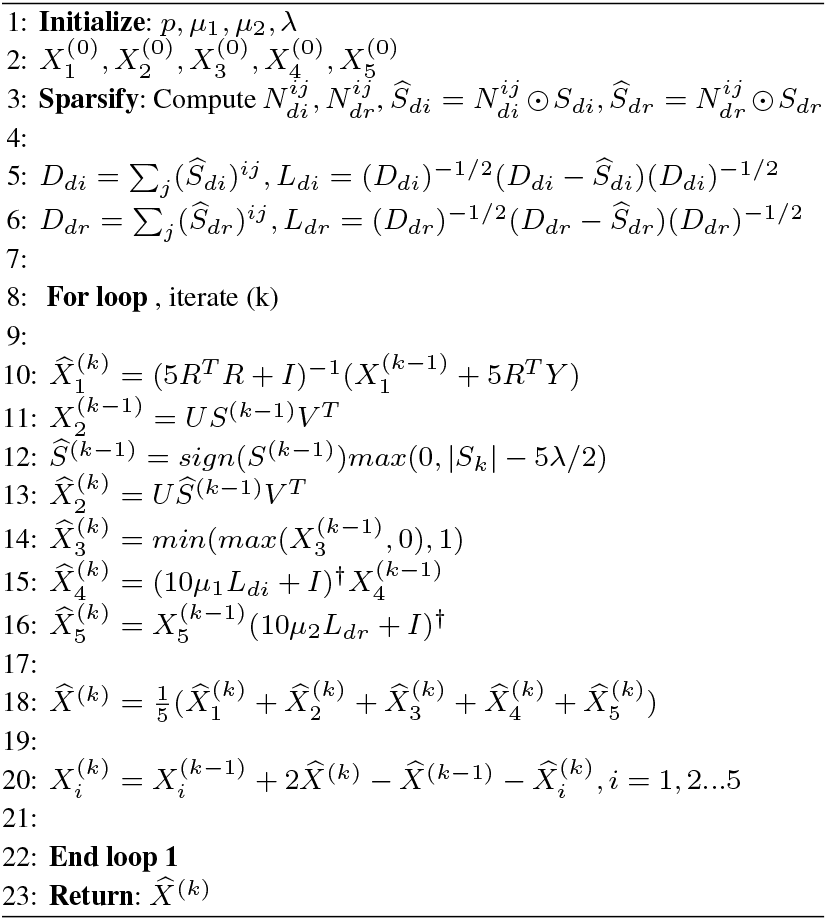

The above procedure is repeated *κ*-times and the average of AUC/AUPR across all the *κ* folds is reported. Figures 4 and 5 show the ROC curves obtained on all the 10 folds after running GR1BMC on both the datasets. The average AUC and AUPR across all the folds has been shown in Tables 2 and 3. As can be observed from the table, GR1BMC performs better than the benchmarks techniques on both the datasets, especially in terms of AUPR. It should be noted that AUPR is a relatively more important metric in this problem since it heavily punishes highly ranked non-associations (false positives), which is crucial given the nature of application as false positive indications would lead to wastage of resources if the proposed indications are tested in clinical experiments.

**Table 2.** Average AUC across 10-fold cross-validation for various techniques while predicting drug-disease associations.

| Datasets | GR1BMC | BNNR | HNRD | DRRS |
| --- | --- | --- | --- | --- |
| Fdataset | 0.9769 | 0.9330 | 0.9420 | 0.9300 |
| Cdataset | 0.9805 | 0.9480 | 0.9500 | 0.9470 |

**Table 3.** Average AUPR across 10-fold cross-validation for various techniques while predicting drug-disease associations.

| Datasets | GR1BMC | BNNR | HNRD | DRRS |
| --- | --- | --- | --- | --- |
| Fdataset | 0.7247 | 0.4410 | 0.5720 | 0.3780 |
| Cdataset | 0.7537 | 0.4710 | 0.6700 | 0.4020 |

**Fig. 4.**
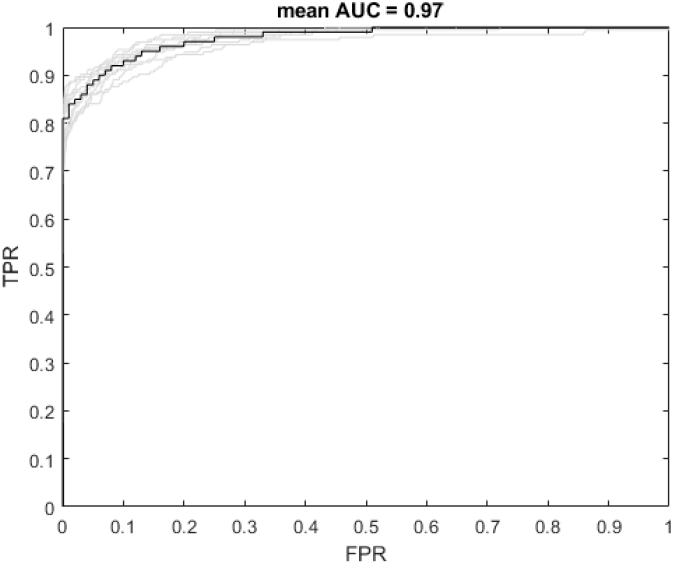
ROC curves obtained for all the 10 folds after applying GR1BMC on Fdataset

**Fig. 5.**
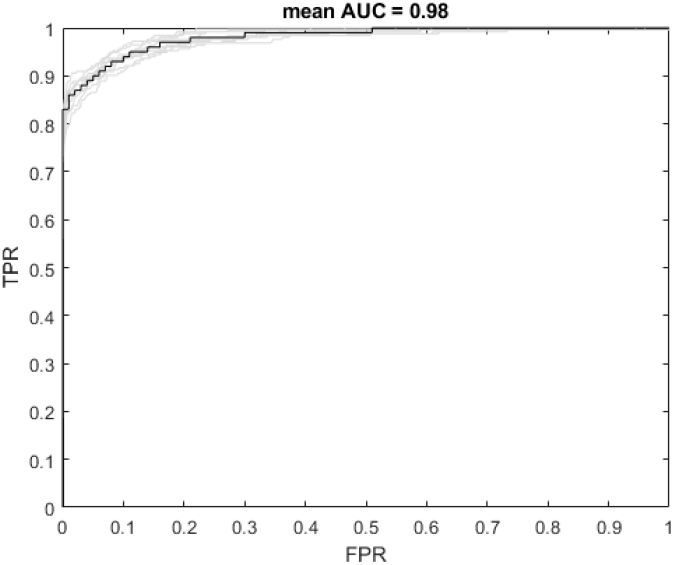
ROC curves obtained for all the 10 folds after applying GR1BMC on Cdataset

### 3.2 Comparison with benchmark techniques

To evaluate the performance of GR1BMC, we compare the results of cross-validation experiments with those of the latest methods proposed for drug-disease association prediction: Bounded nuclear norm regularization (BNNR) Yang *et al.* (2019b), Heterogeneous Network for drug-Disease association prediction (HNRD) Wang *et al.* (2019) and drug repositioning recommendation system (DRRS) Luo *et al.* (2018). BNNR and DRRS are the closest in terms of formulation used to model the problem. Both the methods deploy nuclear norm minimization on a heterogeneous network matrix obtained by integrating drug similarity, disease similarity, association matrix and its transpose; BNNR additionally handles the noise originating from similarities which violate the low-rankness and restrict the predicted values to be in range [0,1]. But, the low-rank property of the heterogeneous matrix is unexplained in both the works; which is a crucial assumption behind nuclear norm minimization. This heterogeneous matrix comprises of associations between drugs and diseases as well as drug-drug and disease-disease similarities. The authors clearly explain validity of the low-rank assumption in association matrix but not for the heterogeneous matrix.

The results of 10-fold cross-validation have been shown in tables 2 and 3. It can be seen that our proposed approach shows competetive performance in terms of area under the ROC curve and is better than the benchmark techniques in temrms of precision and recall also.

### 3.3 Parameter settings

The matrices 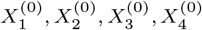 and 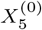 are initialized randomly and the algorithm is run for a fixed number of iterations *κ* (*κ*=20 here) that appears sufficient to reach practical stabilization of the objective function. The running time is in the order of seconds; ppxa takes approximately 4 and 6 seconds on Fdataset and Cdataset respectively. To look for a feasible solution in the space of low-rank association matrices, we need to determine the values of the hyperparameters λ, *μ*_1_ and *μ*_2_. This is done to weigh the importance of nuclear norm term and the trace terms in our objective function for each of the two datasets. The values of *μ*1 and *μ*2 determine the weights given to each of the drug and disease laplacians, hence exhibiting the importance of neighborhood information of drugs and targets in our framework for a dataset. The optimal values of these parameters are found by performing cross validation on the training set and taking the value of parameters from the set {0.01, 0.05, 0.1, 0.5,1, 5, 10}. The values of λ, *μ*_1_ and *μ*_2_ are robust across the datasets and are found to be 0.1,0.05, 0.1 for both the datasets.

### 3.4 Case study to predict novel associations

To asses the practical usage of the proposed algorithm, we perform a case study where we chose 5 candidate drugs to look for their novel indications (dummy drug re-positioning) after prediction the associations using our proposed approach.

We train our model on the known associations on Fdataset. After the matrix completion is done, we rank the remaining candidate diseases for each drug in descending order or predicted association scores.

These rankings or predictions of novel indications for drugs is verified by validating the top-5 indications for any 5 drugs with the public database comparative toxicogenomics database (CTD) Davis *et al.* (2013). We show the validation on the following 5 drugs: Levodopa, Doxorubicin, Amantadine, Flecainide and Metformin.

The indications predicted by GR1BMC and the evidence from CTD is shown in table 4. It can be seen that at least 3 indications are confirmed with the CTD databse for 4 out of 5 drugs and a total of 17 out of 25 predicted associations have evidence in CTD database. Also, the indications which are not verified could be the potential candidates for drug-repositioning and could be explored by medical researchers.

**Table 4.** Top 5 predicted diseases for Levodopa, Doxorubicin, Amantadine, Flecainide and Metformin with their evidence in CTD database

| DRUG INFORMATION |  | DISEASE INFORMATION |  |  |
| --- | --- | --- | --- | --- |
| Drug name | DrugBank ID | Disease name | OMIM ID | Confirmation |
| Levodopa | DB01235 | (PARKINSON DISEASE, LATE-ONSET; PD) | 168600 | CTD confirmed |
|  |  | (DEMENTIA/PARKINSONISM WITH NON-ALZHEIMER AMYLOID PLAQUES) | D125320 | CTD confirmed? |
|  |  | (DYSTONIA 9; DYT9) | D601042 | CTD confirmed |
|  |  | (DEMENTIA, LEWY BODY; DLB) | D127750 |  |
|  |  | (RENAL FAILURE, PROGRESSIVE, WITH HYPERTENSION; RFH1) | D161900 |  |
| Doxorubicin | DB00997 | (COLORECTAL CANCER; CRC) | D114500 | CTD confirmed |
|  |  | (DOHLE BODIES AND LEUKEMIA) | D223350 |  |
|  |  | (RETICULUM CELL SARCOMA) | D267730 | CTD confirmed? |
|  |  | (RENAL CELL CARCINOMA, NONPAPILLARY; RCC) | D144700 | CTD confirmed |
|  |  | (LEUKEMIA, CHRONIC LYMPHOCYTIC, SUSCEPTIBILITY TO, 2) | D109543 | CTD confirmed |
| Amantadine | DB00915 | (PARKINSON DISEASE, LATE-ONSET; PD) | D168600 | CTD confirmed |
|  |  | (DEMENTIA/PARKINSONISM WITH NON-ALZHEIMER AMYLOID PLAQUES) | D125320 | CTD confirmed |
|  |  | (ALZHEIMER DISEASE, FAMILIAL EARLY-ONSET, WITH COEXISTING AMYLOID AND PRION PATHOLOGY) | D605055 | CTD confirmed |
|  |  | (DEMENTIA, LEWY BODY; DLB) | D127750 | CTD confirmed |
|  |  | (ALZHEIMER DISEASE; AD) | D104300 | CTD confirmed |
| Flecainide | DB01195 | (ATRIAL FIBRILLATION, FAMILIAL, 1; ATFB1) | D608583 | CTD confirmed |
|  |  | (HYPERTENSION, DIASTOLIC, RESISTANCE TO) | D608622 | CTD confirmed |
|  |  | (RENAL FAILURE, PROGRESSIVE, WITH HYPERTENSION; RFH1) | D161900 |  |
|  |  | (INSENSITIVITY TO PAIN WITH HYPERPLASTIC MYELINOPATHY) | D147530 |  |
|  |  | (STROKE, ISCHEMIC) | D601367 |  |
| Metformin | DB00331 | (DIABETES MELLITUS, INSULIN-DEPENDENT, 2) | D125852 | CTD confirmed |
|  |  | (COLORECTAL CANCER; CRC) | D114500 | CTD confirmed |
|  |  | (HYPERLIPOPROTEINEMIA, TYPE V) | D144650 | CTD confirmed |
|  |  | (ENDOMETRIOSIS, SUSCEPTIBILITY TO, 1) | D131200 |  |
|  |  | (UTERINE ANOMALIES) | D192000 |  |

## 4 Conclusion

The huge amount of time and efforts taken for the development drugs calls for the need for efficient and reliable computational methods to assist drug re-positioning. In this paper, we present a novel approach to predict drug-disease indications based on parallel proximal algorithm, which benefits from guaranteed convergence and great numerical performance. Cross validation and experiments on gold standard dataset demonstrate the superiority of the proposed approach over the benchmark techniques. The practical usage is also validated by the case study where novel indications for existing drugs are found and majority are validated with the CTD database. The proposed method is generic and can be applied to other association/interaction prediction problems such as proteinprotein interaction prediction, human microbe-disease association (MDA) prediction, gene-disease association prediction, etc.

## Acknowledgements

This manuscript has been submitted to the preprint server-bioRxiv

## Footnotes

1 The code of GR1BMC is available at https://github.com/aanchalMongia/GROBMC-PPXA-DDA

## References

Belkin, M. et al. (2006). Manifold regularization: A geometric framework for learning from labeled and unlabeled examples. Journal of machine learning research, 7(Nov), 2399–2434.

Candès, E. J. and Recht, B. (2009). Exact matrix completion via convex optimization. Foundations of Computational mathematics, 9(6), 717.

Cherni, A. et al. (2017). Palma, an improved algorithm for dosy signal processing. Analyst, 142(5), 772–779.

Chiang, A. P. and Butte, A. J. (2009). Systematic evaluation of drug-disease relationships to identify leads for novel drug uses. Clinical Pharmacology & Therapeutics, 86(5), 507–510.

Combettes, P. L. and Pesquet, J.-C. (2011). Proximal splitting methods in signal processing. In Fixed-point algorithms for inverse problems in science and engineering, pages 185–212. Springer.

Dai, W. et al. (2015). Matrix factorization-based prediction of novel drug indications by integrating genomic space. Computational and mathematical methods in medicine, 2015.

Das, S. et al. (2019). Repurposing disulfiram to target infections caused by non-tuberculous mycobacteria. Journal of Antimicrobial Chemotherapy, 74(5), 1317–1322.

Davis, A. P. et al. (2013). The comparative toxicogenomics database: update 2013. Nucleic acids research, 41(D1), D1104–D1114.

Gottlieb, A. et al. (2011). Predict: a method for inferring novel drug indications with application to personalized medicine. Molecular systems biology, 7(1).

Hamosh, A. et al. (2002). Online mendelian inheritance in man (omim), a knowledgebase of human genes and genetic disorders. Nucleic acids research, 30(1), 52–55.

He, S. et al. (2015). Repurposing of the antihistamine chlorcyclizine and related compounds for treatment of hepatitis c virus infection. Sciencetranslational medicine, 7(282), 282ra49–282ra49.

Hu, G. and Agarwal, P. (2009). Human disease-drug network based on genomic expression profiles. PloS one, 4(8).

Jadamba, E. and Shin, M. (2016). A systematic framework for drug repositioning from integrated omics and drug phenotype profiles using pathway-drug network. BioMed research international, 2016.

Jiang, H.-J. et al. (2019). Predicting drug-disease associations via using gaussian interaction profile and kernel-based autoencoder. BioMed research international, 2019.

Kaul, G. et al. (2019). Update on drug-repurposing: is it useful for tackling antimicrobial resistance?

Lamb, J. et al. (2006). The connectivity map: using gene-expression signatures to connect small molecules, genes, and disease. science, 313(5795), 1929–1935.

Luo, H. et al. (2016). Drug repositioning based on comprehensive similarity measures and bi-random walk algorithm. Bioinformatics, 32(17), 2664–2671.

Luo, H. et al. (2018). Computational drug repositioning using low-rank matrix approximation and randomized algorithms. Bioinformatics, 34(11), 1904–1912.

Martinez, V. et al. (2015). Drugnet: Network-based drug-disease prioritization by integrating heterogeneous data. Artificial intelligence in medicine, 63(1), 41–49.

Pustelnik, N. et al. (2011). Parallel proximal algorithm for image restoration using hybrid regularization. IEEE transactions on Image Processing, 20(9), 2450–2462.

Steinbeck, C. et al. (2003). The chemistry development kit (cdk): An open-source java library for chemo-and bioinformatics. Journal of chemical information and computer sciences, 43(2), 493–500.

Tanimoto, T. T. (1958). An elementary mathematical theory of classification and prediction. 1958. International Business Machines Corporation.

Thakare, R. et al. (2019). Repurposing disulfiram for treatment of staphylococcus aureus infections. International journal of antimicrobial agents, 53(6), 709–715.

Van Driel, M. A. et al. (2006). A text-mining analysis of the human phenome. European journal of human genetics, 14(5), 535–542.

Villanueva-Lozano, H. et al. (2018). Clinical evaluation of the antifungal effect of sertraline in the treatment of cryptococcal meningitis in hiv patients: a single mexican center experience. Infection, 46(1), 25–30.

Walters, W. P. et al. (2011). What do medicinal chemists actually make? a 50-year retrospective. Journal of medicinal chemistry, 54(19), 6405–6416.

Wang, W. et al. (2014). Drug repositioning by integrating target information through a heterogeneous network model. Bioinformatics, 30(20), 2923–2930.

Wang, Y. et al. (2013). Drug repositioning by kernel-based integration of molecular structure, molecular activity, and phenotype data. PloS one, 8(11).

Wang, Y. et al. (2019). Drug-disease association prediction based on neighborhood information aggregation in neural networks. IEEE Access, 7, 50581–50587.

Wishart, D. S. et al. (2006). Drugbank: a comprehensive resource for in silico drug discovery and exploration. Nucleic acids research, 34(suppl_1), D668–D672.

Xuan, P. et al. (2019). Heterodualnet: A dual convolutional neural network with heterogeneous layers for drug-disease association prediction via chou’s five-step rule. Frontiers in pharmacology, 10.

Yang, J. et al. (2014). Drug-disease association and drug-repositioning predictions in complex diseases using causal inference-probabilistic matrix factorization. Journal of chemical information and modeling, 54(9), 2562–2569.

Yang, K. et al. (2019a). Predicting drug-disease associations with heterogeneous network embedding. Chaos: An Interdisciplinary Journal of Nonlinear Science, 29(12), 123109.

Yang, M. et al. (2019b). Drug repositioning based on bounded nuclear norm regularization. Bioinformatics, 35(14), i455–i463.

Yu, L. et al. (2016). Prediction of new drug indications based on clinical data and network modularity. Scientific reports, 6, 32530.

Zhang, W. et al. (2018). Predicting drug-disease associations by using similarity constrained matrix factorization. BMC bioinformatics, 19(1), 1–12.

